# Effect of mutations in the SARS-CoV-2 spike protein on protein stability, cleavage, and cell-cell fusion function

**DOI:** 10.1101/2021.01.24.428007

**Authors:** Chelsea T. Barrett, Hadley E. Neal, Kearstin Edmonds, Carole L. Moncman, Rachel Thompson, Jean M. Branttie, Kerri Beth Boggs, Cheng-Yu Wu, Daisy W. Leung, Rebecca E. Dutch

## Abstract

The SARS-CoV-2 spike protein (S) is the sole viral protein responsible for both viral binding to a host cell and the membrane fusion event needed for cell entry. In addition to facilitating fusion needed for viral entry, S can also drive cell-cell fusion, a pathogenic effect observed in the lungs of SARS-CoV-2 infected patients. While several studies have investigated S requirements involved in viral particle entry, examination of S stability and factors involved in S cell-cell fusion remain limited. We demonstrate that S must be processed at the S1/S2 border in order to mediate cell-cell fusion, and that mutations at potential cleavage sites within the S2 subunit alter S processing at the S1/S2 border, thus preventing cell-cell fusion. We also identify residues within the internal fusion peptide and the cytoplasmic tail that modulate S cell-cell fusion. Additionally, we examine S stability and protein cleavage kinetics in a variety of mammalian cell lines, including a bat cell line related to the likely reservoir species for SARS-CoV-2, and provide evidence that proteolytic processing alters the stability of the S trimer. This work therefore offers insight into S stability, proteolytic processing, and factors that mediate S cell-cell fusion, all of which help give a more comprehensive understanding of this highly sought-after therapeutic target.

## Introduction

Severe acute respiratory syndrome coronavirus 2 (SARS-CoV-2) is the causative viral agent of the ongoing coronavirus disease of 2019 (COVID-19) global pandemic. Thus far, COVID-19 has impacted over 86 million people globally, resulting in the death of more than one and a half million individuals [1]. Due to the widespread global impact of this pandemic, a concerted effort has been made to rapidly develop a vaccine or antiviral treatment.

The SARS-CoV-2 spike (S) protein is the major transmembrane glycoprotein studding the surface of the viral particle, and is exclusively responsible for viral attachment and cell entry, thus making it the major target of current vaccine strategies and antiviral therapeutics [2]. The S protein consists of two distinct subunits: the S1 subunit, which binds to the known host receptor, angiotensin converting enzyme 2 (ACE2) [3-11], and the S2 subunit that promotes the viral-to-host cell membrane fusion event needed for viral infection [2, 8, 12-18]. Most known coronavirus (CoV) S proteins undergo two post-translational proteolytic cleavage events, one at the border of the S1 and S2 subunits, and one downstream within the S2 subunit (termed S2’) [2, 13, 15-21].

Similar to several other CoVs, SARS-CoV-2 likely utilizes bats as a reservoir species, specifically *Rhinolophus affinis* or horseshoe bats [11, 22-25]. SARS-CoV-2 has 96% sequence identity to a CoV found in this bat population, RaTG13, with limited differences between them [25]. One difference is the polybasic, PRRA, insertion at the S1/S2 border which gives this site the canonical sequence requirements for cleavage by the cellular proprotein convertase furin [26-29]. This change may be a key factor in the zoonotic transmission of SARS-CoV-2. The presence of a furin consensus sequence at the cleavage site has been observed in other human infecting CoVs [26, 30-32], including highly pathogenic forms of influenza [33, 34] and previous studies have demonstrated its functional significance. For SARS-CoV-2, the insertion is suggested to allow for expanded cellular tropism and infectivity [13, 26, 35, 36]. For most CoVs, cleavage at a downstream S2’ site may be carried out by a number of cellular proteases, including serine proteases like transmembrane serine protease 2 (TMPRSS2), or endopeptidases, including members of the cathepsin family [13, 14, 19-21].

Following receptor binding by the S1 subunit and priming by proteolytic cleavage, the S2 subunit of S promotes the critical membrane fusion step of viral entry by undergoing dynamic conformational changes to promote merging of the viral and host cell membranes [10, 35, 37]. For entry of SARS-CoV-2, cleavage at the S1/S2 border (by furin or a similar protease), is critical for TMPRSS2 cleavage and entry at the plasma membrane. However, when S1/S2 border cleavage is blocked, viral entry can be mediated through endosomal compartments with proteolytic cleavage carried out by a member of the cathepsin family, similar to the entry pathway of SARS-CoV [10, 35, 37-39]. In addition to promoting virus-cell fusion during viral particle entry, S can also promote cell-cell fusion, a pathogenic effect observed in the lungs of COVID-19 patients where neighboring cells fuse together to form large multi-nucleated cells, termed syncytia [40-45]. While the role of cellular proteases and S cleavage in viral entry is being extensively investigated, insight into the cleavage requirements for cell-cell fusion in SARS-CoV-2 remains more limited. Recent studies have suggested that S cleavage at the S1/S2 border is critical for cell-cell fusion, and TMPRSS2, while not required, appears to enhance this cell-cell fusion [37, 40, 46, 47]. However, relatively little is known about the timing and efficiency of these cleavage events, and how mutations in S may affect the process.

Though CoVs mutate at a slower rate than most RNA viruses due to the presence of viral proofreading machinery, a meta-analysis of genomes of SARS-CoV-2 strains found several mutations within S circulating in significant percentages of the analyzed populations [48, 49]. The most common mutation, now found in most of the global population, is an aspartate to glycine mutation at residue 614 (D614G) in the S1 subunit. Additional mutations throughout the S1 and S2 subunits of S have been found in a smaller percentage of the viral population. Since S2 contains the fusion machinery, mutations in this region may have an impact on overall protein stability and fusion. Understanding the effects of mutations in this region will allow for a more comprehensive understanding of the overall S function.

We tested wild-type (wt) SARS-CoV-2 S and variants in different host cell strains to analyze the effects on stability, proteolytic processing, and cell-cell fusion. Here we demonstrate that furin cleavage of S at the S1/S2 border is required for efficient cell-cell fusion, and that the presence of TMPRSS2 in target cells enhances S mediated cell-cell fusion, consistent with previous studies [37, 46]. We also show that mutations of the cleavage sites at the S1/S2 border, S2’ site, or a cathepsin L (cath L) cleavage site, conserved from SARS-CoV S, all reduce initial cleavage at the S1/S2 border during viral protein synthesis, suggesting that mutations downstream of the S1/S2 border likely alter the overall conformation of the protein. Additionally, we identify two S2 subunit residues, one in the internal fusion peptide and another in the cytoplasmic tail, that alter protein fusion function when mutated without changing overall protein expression and cleavage, providing more insight into regions of the protein important for the regulation of the fusion process. Finally, we demonstrate protein turnover and cleavage kinetics in a range of host cells, as well as in the presence of several exogenous proteases, providing a more comprehensive picture of the S protein.

## Results

### Stability and proteolytic cleavage of SARS-CoV-2 Spike in Various Cell Lines

To examine the stability and cleavage patterns of SARS-CoV-2 S in a range of mammalian cell lines, the following cells were transiently transfected with pCAGGS-S: Vero, A549, MEFs, Cath L-MEFS, and LoVo cells (a human colon carcinoma line that does not express functional furin). Stability of S and the timing of proteolytic processing were determined by pulse-chase labeling and immunoprecipitation. S protein detected from immunoprecipitation was observed as two bands, a band around 150 kDa corresponding to an un-cleaved full-length species of the protein, labeled S, and a band around 97 kDa corresponding to a species of S cleaved at the border of the S1 and S2 subunits, labeled S2 (Fig. 1a). After a one-hour chase, a band corresponding to S2 was observed in Vero, A549, and both MEF cell lines (Fig. 1a). In LoVo cells, a band corresponding to the S2 subunit did not appear until four hours of chase, verifying that lack of furin impedes efficient processing at S1/S2, and that the S1/S2 border can be cleaved by cellular protease other than furin (Fig. 1a) in a slower and less efficient process. Veros, A549s, MEFs, and Cath L-MEFs displayed similar cleavage patterns over time, while LoVo cells displayed significantly less cleavage at two and four hours. LoVo cells had only 2% cleavage at two hours and 18% cleavage at four hours, compared to about 20-40% at two hours and 30-60% at four hours for all other cell types (p<0.05). However, LoVo cells reached cleavage levels similar to the other cell lines at later chase time points (Fig. 1b). Bands smaller than 90 kDa that would correspond to cleavage at the S2’ site were not observed in any cell line. In the examined cell lines, expressed S remained stable through the first four hours (Fig. 1c). By 24 hours post label, only 20-30% of the original labeled protein remained for all cell lines.

**Figure 1:**
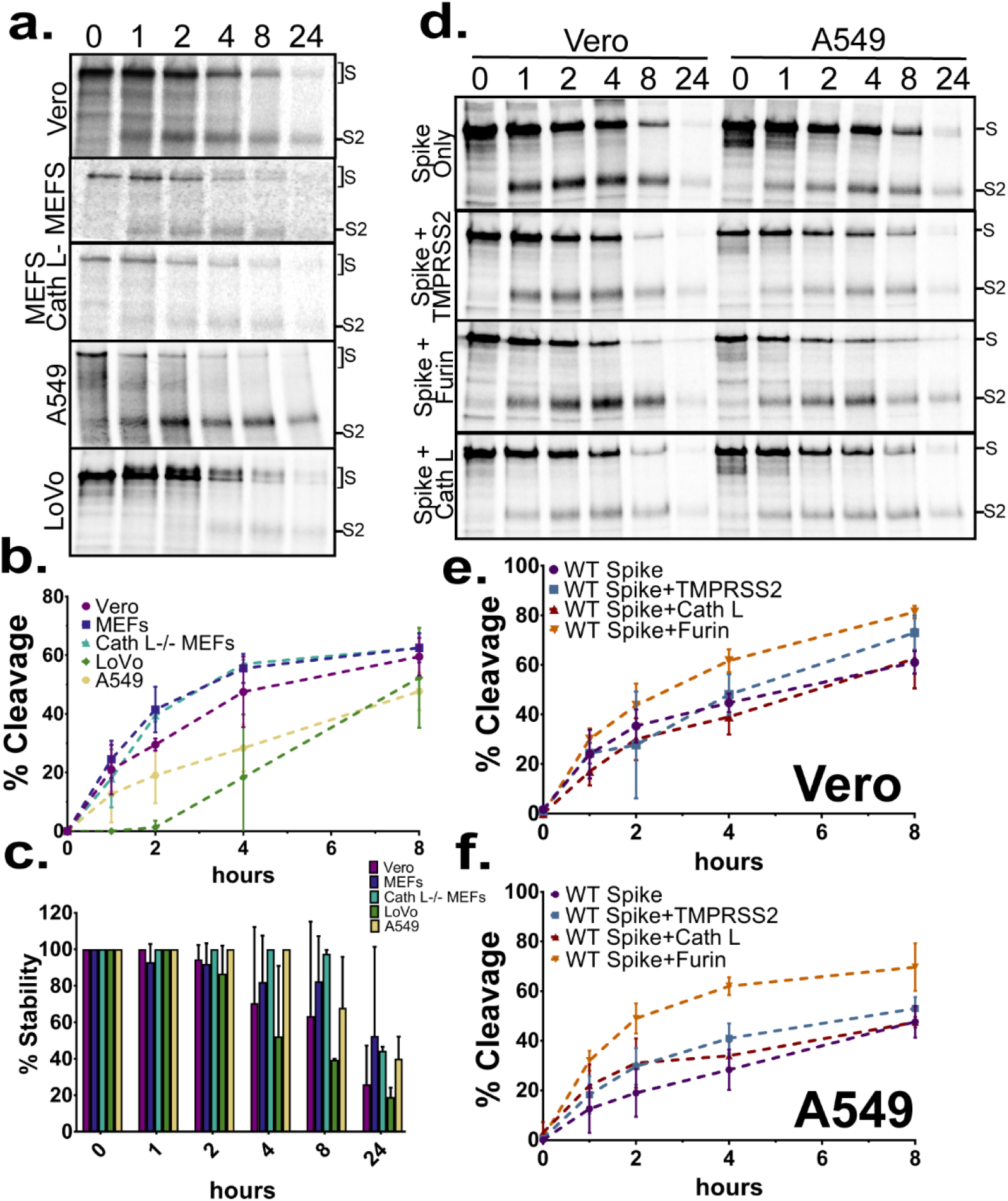
SARS-CoV-2 Spike is cleaved at the S1/S2 subunit border in a variety of cell lines. a) The indicated cell types transiently expressing S were metabolically labeled for one hour, and chased for times indicated (hours). Band densitometry was used to quantify bands representing full length S or S cleaved at the S1/S2 border (S2) (b) Percent cleavage [S2 divided by S plus S2] and (b) Overall protein stability [Total S, S plus S2, for each time point, normalized to time point 0] were calculated for spike in each cell line (n=3). d) S alone, or S with proteases transiently expressed in Vero and A549 cells, cells were metabolically labeled, and chased for the times indicated (hours). Percent cleavage was measured using band densitometry in both (e) Vero and (f) A549 cells (b, c, e, f are represented as the average ± SD for 3 independent experiments).

Several studies have examined the cellular proteases involved in the cleavage of S. Furin and TMPRSS2 appear to play key roles in cleavage at the S1/S2 border and S2’ site, respectively [26, 35, 50-52]. Additionally, lysosomal proteases such as cath L/B can be utilized for viral entry in TMPRSS2 deficient cells [10, 38, 46]. To examine how higher expression levels of these proteases affect S stability and cleavage, Vero and A549 cells were transiently transfected with S alone or S with TMPRSS2, furin, or cath L. Pulse-chase analysis demonstrated that the transient expression of TMPRSS2 or cath L did not affect the cleavage pattern of S (Fig. 1d and 1e, S1b), and a band corresponding to S2’ cleavage was not observed in either Veros or A549s. However, transient over-expression of furin increased the cleavage observed at the S1/S2 border in Veros at four and eight hours of chase (p<0.05) and at all times after zero for A549s (p<0.01 for one- and eight-hour chase, p<0.0001 for two- and four-hour chase times) (Fig. 1e and 1f). This suggests that the normal levels of cellular furin can eventually promote maximal levels of S1/S2 cleavage in both Veros and A549s, but over-expression of furin facilitates more rapid cleavage of the S1/S2 border. Interestingly, in both experiments (Fig. 1a and 1d) some un-cleaved S remains even after 24 hours, indicating that a small portion of the S population is not cleaved by furin or other endogenous proteases in these cell lines. Finally, overall protein stability was not affected by co-expression of any tested proteases (Fig. S1b).

### Spike Mediated Cell-Cell Fusion

The S2 subunit of S mediates both viral-cell fusion and cell-cell fusion [40-42], with cell-cell fusion readily observed both in a laboratory setting and in the lungs of SARS-CoV-2 infected patients [40-45]. To better understand the requirements and contribution of cellular proteases to S2 mediated cell-cell fusion, we performed syncytia and reporter gene assays. For syncytia analysis, a small number of syncytia, were observed at 24 hpt in all samples (Fig. 2a). At 48 hpt, similar numbers of large syncytia were observed with S alone or S co-expressed with TMPRSS2 or cath L (Fig. 2b). However, co-expression of S with furin resulted in increased syncytia formation. The cells exhibited nearly complete fusion, suggesting that the presence of exogenous furin further increases S mediated cell-cell fusion (Fig. 2b, panel 3).

**Figure 2:**
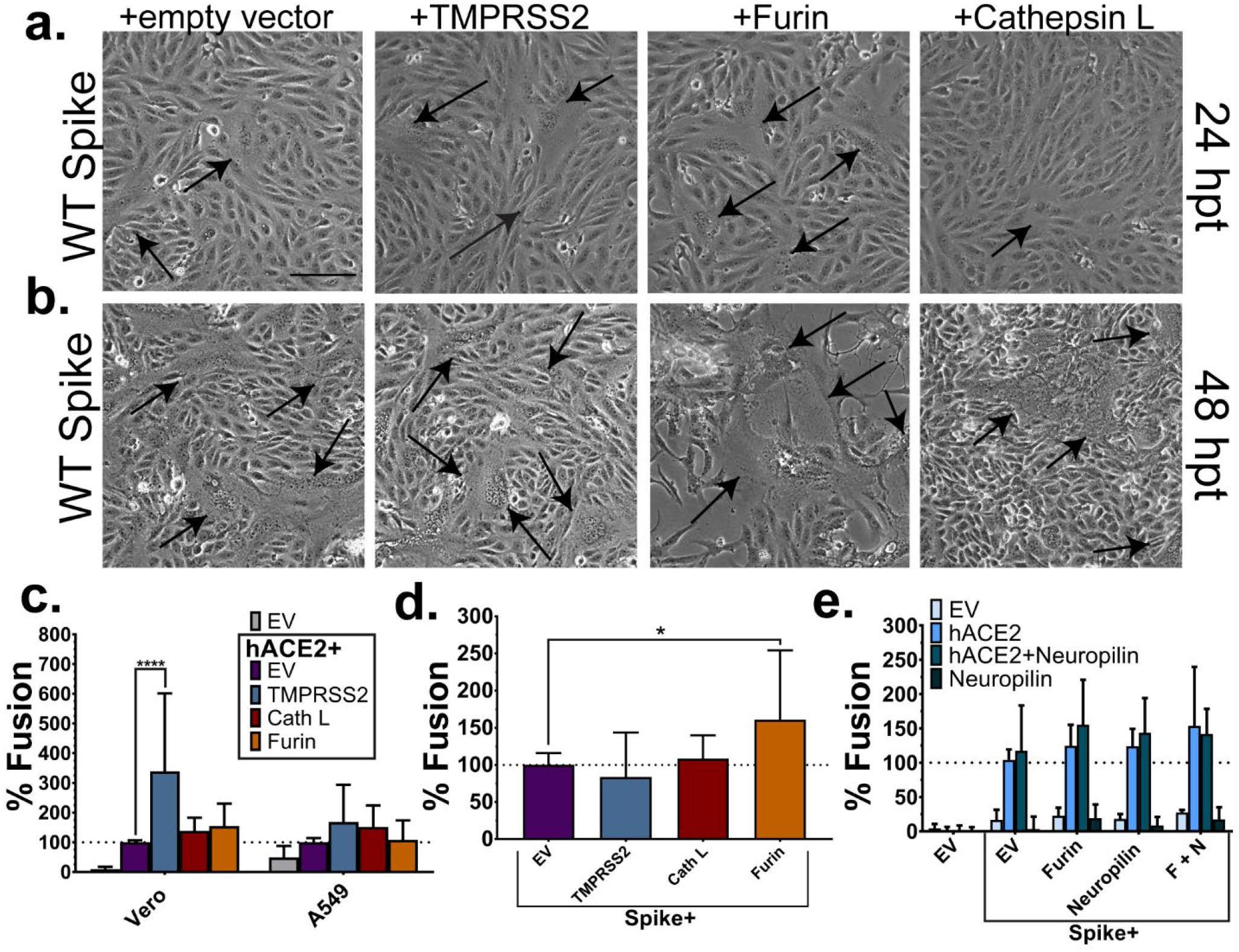
CoV-2 spike alone mediates cell-cell fusion. Veros expressing S and TMPRRS2, furin, or cathepsin L were imaged at 24 (a) and 48 (b) hpt for syncytia formation (black arrows). Magnification bar is 100μM. c) A luciferase reporter gene assay was performed with target cells (BSR/T7s expressing hACE2 and additional proteases) overlaid onto effector cells (Vero or A549s expressing S) for 9 hours. d) Luciferase reporter gene experiment was performed with additional proteases co-expressed with S in Veros and overlain with target cells expressing hACE2. e) The effect of Neuropilin in both target and effector (Vero) cells was examined with a luciferase reporter gene assay. Effector cells expression is listed along the x-axis. Target cell expression is listed in the graph legend. Results expressed as the percent fusion normalized to samples with S in the effector cells, and hACE2 only in the target cells (c-e are average ± SD for 3 independent experiments, performed in duplicate). Significance was determined by two-way ANOVA. *: p < 0.05, ****: p<0.0001

To quantitate S mediated cell-cell fusion, luciferase reporter gene fusion assays were performed (Fig. S2a), using a nine hour overlay that was determined to be optimal (Fig. S2b). To characterize the role of cellular proteases in the hACE2 expressing target cells, S-expressing effector cells were overlaid with target cells containing hACE2 alone or hACE2 with TMPRSS2, furin, or cath L. The amount of plasmid transfected was kept constant by supplementing with a plasmid encoding an empty expression vector (EV). When Vero cells were used as the S-expressing effector cell and TMPRSS2 was present in the target cells, a significant increase in fusion was observed. This is consistent with the concept that TMPRSS2 plays a role in fusion after or during the hACE2 (receptor) binding step in the fusion cascade (Fig. 2c) [10, 12, 32, 37, 46], although the presence of TMPRSS2 in these target cells also appeared to process hACE2 (Fig. S2c, also observed in[40]). In samples with cath L or furin in the target cells, fusion levels were similar to hACE2+EV (Fig. 2c). When A549 cells were used as the S-expressing effector cell, none of the conditions produced statistically significant differences from background levels (Fig. 2c), so Vero cells were used as the effector cells for the remainder of the experiments performed.

Having analyzed the function of proteases in the target cells, we were also interested in the role of proteases present in the S-expressing effector cells. To test this, EV, TMPRSS2, cath L, or furin were co-expressed with S and samples were overlaid with target cells expressing hACE2. Similar to what we observed in syncytia assays, only co-expression of S and furin produced a statistically significant increase in fusion. This increase is likely due to the increase in the amount of cleaved protein present when S is co-expressed with furin (Fig. 1e).

Neuropilin-1 has been suggested as a co-receptor for SARS-CoV-2 S and may be important for the viral infection infiltrating the neuronal network [53-55]. To assess the contribution of neuropilin in cell-cell fusion, effector cells were transfected with S and either EV, furin, neuropilin, or furin and neuropilin (F+N). Target cells were transfected with EV, hACE2, neuropilin, or hACE2 and neuropilin. However, no significant increase in fusion was observed when neuropilin was present in either the target or effector cells (Fig. 2e), suggesting that neuropilin does not appear to play a significant role in cell-cell mediated fusion. Interestingly, when neuropilin is co-expressed in S containing effector cells, there is no difference observed in fusion compared to samples with S+EV, suggesting that neuropilin also does not have an inhibitory effect (Fig. 2e). Additionally, when neuropilin alone is expressed in the target cells, fusion levels above background levels are not observed. This indicates that in cell-cell fusion, S binding hACE2 appears to be the major interaction during the receptor attachment function.

### Importance of CoV-2 cleavage sites

Early protein sequence analysis of CoV-2 S protein demonstrated the presence of three potential cleavage sites [26]: a putative furin cleavage site at the S1/S2 border; a conserved site 10 residues downstream from the S1/S2 border, shown to be cleaved by cath L in SARS-CoV; and the S2’ site which is potentially cleaved by TMPRSS2 [26]. To functionally understand the role of each cleavage site in S cell-cell fusion, a series of mutants were made. Alanine mutations of all the residues within each potential cleavage site (S1/S2 AAAAA, Cath L AAAA, S2’ AA), and single alanine mutations at the terminal arginine of the S1/S2 border and S2’ site (S1/S2 PRRAA, S2’ KA) were created. Finally, a mutant with residues (PRRA) upstream of the S1/S2 border deleted (del. PRRA), leaving a single R residue at this site, was made, creating an S1/S2 border similar to SARS-CoV S (Fig. 3A). Pulse-chase analysis (Fig. 3b) showed that all mutants had similar protein turnover compared to wt S in Veros. However, in A549s several mutants demonstrated more rapid protein turnover than wt S at later chase time points. Surprisingly, mutations at all three sites led to either a complete loss or significant delay in the proteolytic processing of the S protein at the S1/S2 border, indicated by the lack of a band corresponding to the S2 subunit. This suggests that mutations at distal sites can strongly influence cleavage at S1/S2. After an eight-hour chase, no cleavage at the S1/S2 border was observed for the mutants del. PRRA and S1/S2 AAAAA, confirming that deletion or mutation of the furin consensus prevents cleavage at this site. For all other mutants, cleavage at the S1/S2 border reached 30-50% of wt levels in both Vero and A549 cells the eight-hour time point (Fig. 3c and 3d). Accurate analysis of protein cleavage was not possible by the 24-hour time point, since only 20-30% of protein remained (Fig. S1b). Finally, surface biotinylation showed that both total and cell surface expression of all mutants were similar to wt S levels (Fig. 3e, f, and g).

**Figure 3:**
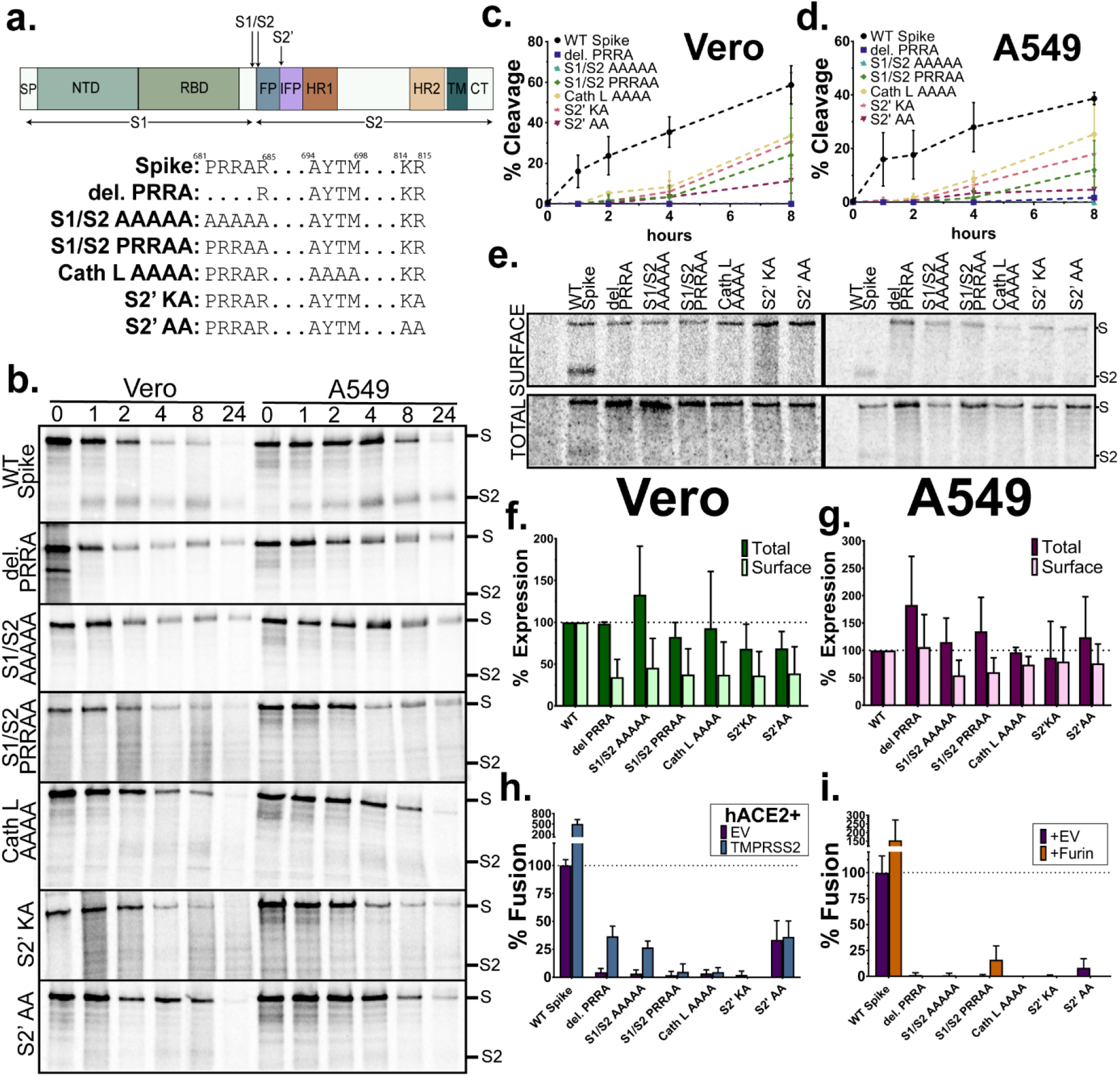
Mutations at all three potential spike cleavage sites reduce cleavage at the S1/S2 subunit border. a) Full or partial alanine substitution mutations were made at each of the three potential cleavage sites. b) Plasmids expressing wt S or mutants were transfected into Veros and A549s, cells were metabolically labeled for one hour, and chased for the times indicated. Percent cleavage was determined in (c) Veros and (d) A549s (average ± SD for 3 independent experiments) e) Surface biotinylation was performed on cells expressing wt S and each mutant. Cells were radiolabeled for 6 hours. Protein expression in (f) Vero and (g) A549 cells, results are normalized to wt S, and error bars represent the standard deviation (average ± SD for 3 independent experiments). h) A luciferase reporter gene assay was performed using target cells expressing hACE2 and EV or TMPRRSS2, and effector (Vero) cells with wt S or each mutant. i) Luciferase reporter gene analysis with cells expressing hACE2 and effector (Vero) cells transfected with S or S mutants and EV or furin expressing plasmids. Results of both reporter gene assays are shown normalized to samples with wt S in the effector with hACE2 in target cells (average ± SD for 3 independent experiments, performed in duplicate).

To assess the effects of the mutations on cell-cell fusion, syncytia formation assays in Vero cells were performed. While syncytia were readily observed in all samples containing wt S, none of the mutants exhibited syncytia formation at 24 or 48 hpt when expressed alone (Fig. S3, panel 2). Addition of TMPRSS2 did not recover syncytia formation in any mutant (Fig. S3, panel 3), and the addition of furin only recovered syncytia formation in the S1/S2 PRRAA mutant (Fig. S3, panel 4, syncytia denoted with black arrows). To analyze this result, cells were lysed following the 48-hour imaging and protein levels examined by western blot. Results showed that co-expression of furin with the S1/S2 PRRAA mutant restored cleavage at the S1/S2 border, while all other mutants did not show cleavage at this site (data not shown). This suggests that cleavage at the S1/S2 border is critical for cell-cell fusion, and that the double R motif in the PRRAA mutant can be cleaved by over-expressed furin.

Luciferase reporter gene analysis of fusion in Veros transfected with wt S or each mutant showed similar results to the syncytia assays, with none of the mutants showing fusion levels above background (Fig. 3h). Interestingly, the S2’ AA mutant displayed high background levels, suggesting this mutant may have a conformational change, or characteristics that increase receptor binding or alter S2 trimeric association, leading to higher background signals. Reporter gene assays were also carried out with addition of transiently expressed furin in the S-expressing effector cells, but no significant increases in fusion were observed. Since all cleavage mutants created reduced cleavage at the S1/S2 subunit border, the reductions in cell-cell fusion may be attributable to loss of cleavage at this site.

### Effect of Circulating S Mutations on Protein Stability, Cleavage, and Fusion

An early examination revealed several mutations in the S protein gene in circulating viral strains [48, 49], including the D614G substitution now found in most of the global SARS-CoV-2 strains [48, 56-62]. The D614G mutation lies in the S1 subunit of the protein, just downstream of the receptor binding domain, and is proposed to play a critical role in receptor binding by alteration of the positioning of the receptor binding domain. Other mutations in circulating strains were found throughout the S2 subunit [49]. To assess the effect(s) of these mutations, we created the mutants D614G, A831V, D839Y/N/E, S943P, and P1263L (Fig. 4a). Pulse-chase analysis in Veros and A549s (Fig. 4b, c) demonstrated that all circulating mutants tested exhibited protein turnover at similar rates as wt S in both cell lines (Fig. S1d). Surface biotinylation confirmed that all tested mutants displayed total protein and surface protein levels comparable to wt S, suggesting that none of the mutants caused major defects or enhancement of protein trafficking to the cell surface (Fig. 4d, e). Syncytia formation and evaluation of protein location by immunofluorescence were similar between all mutants and wt S (Fig. S4). Interestingly, cellular extensions containing the S protein were observed for the wt and each of the mutants (Fig. S4, white arrows) [63]. Finally, luciferase reporter gene assays were performed. While most of the mutants displayed fusion levels similar to wt S, three mutants exhibited significant changes (Fig. 4f). D839Y and D839N displayed significantly reduced levels of fusion compared to wt (p<0.01 and p<0.05, respectively), and P1263L showed a significant increase in fusion compared to wt (p<0.05). These changes in fusion cannot be attributed to differences in cell surface protein expression or cleavage levels, suggesting that residues near the internal fusion peptide, where D839 is located, and residues in the cytoplasmic tail, where P1263 is located, may play an important role in controlling the fusion cascade.

**Figure 4:**
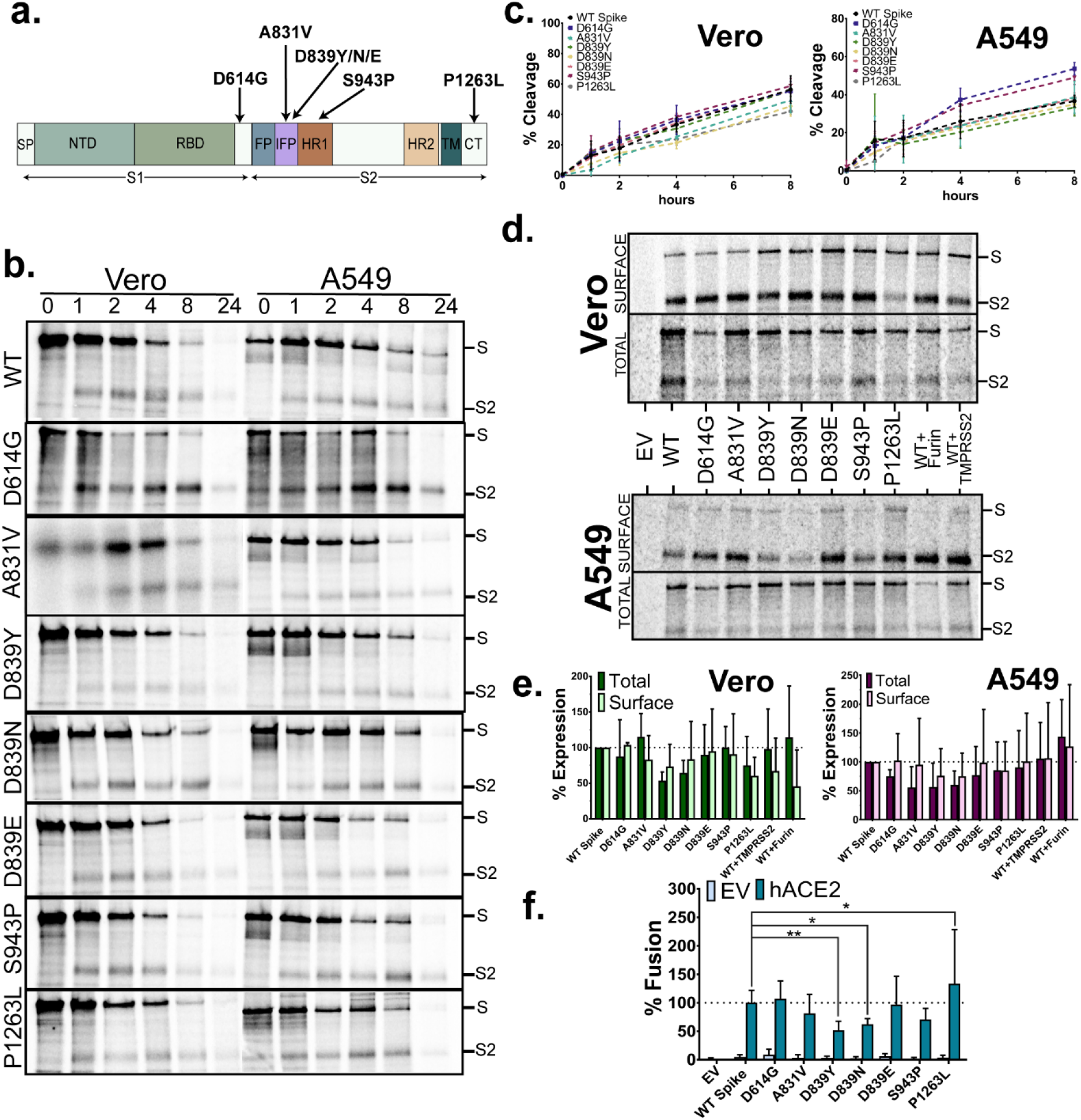
Spike S2 subunit mutations found in circulating strains variably affect spike mediated cell-cell fusion. a) Mutations in the S2 subunit of S identified in circulating SARS-CoV-2 strains, b) Wt S or the mutants were transfected into Veros and A549s, metabolically labeled for one hour, and chased for the times indicated. Percent cleavage was determined in (c) Veros and A549s (average ± SD for 3 independent experiments). d) Surface biotinylation on cells expressing wt S or each mutant. e) Total and surface protein expression normalized to wt S (average ± SD for 3 independent experiments). f) A luciferase reporter gene assay was performed using target cells expressing EV or hACE2, overlaid onto effector cells transfected with wt S or each mutant. Results are normalized to samples with wt S in the effector cells and hACE2 in target cells (average ± SD for 3 independent experiments, performed in duplicate). Significance was determined by two-way ANOVA, *: p<0.05, **: p<0.01.

### Trypsin accessibility and protein-protein association in select Spike mutants

Since all the S cleavage site mutants exhibited defects in cleavage at the S1/S2 border, we evaluated the accessibility of this site using a trypsin treatment assay to determine if the lack of cleavage was due to misfolding in the S1/S2 border region. Veros or A549s were transfected with wt S or each cleavage mutant and metabolically labeled. Cell surface proteins were biotinylated and then cells were either left untreated or treated with 0.3 µg/μl of TPCK-Trypsin prior to lysis. When treated with exogenous TPCK-Trypsin, both the del. PRRA and S1/S2 PRRAA mutants were efficiently cleaved at the S1/S2 border, shown by the appearance of a band corresponding to S2 in the lanes treated with trypsin (Fig. 5a, quantified in Fig. 5b). This suggests that the observed defects in cleavage at the S1/S2 border are not due to inaccessibility at the site, but rather to the removal of the furin consensus sequence. Interestingly, mutations at the downstream cath L or S2’ potential cleavage sites also render defects in protein cleavage at the S1/S2 border site. However, treatment with exogenous trypsin did not significantly affect the amount of cleavage observed, a result consistent with a change in conformation that renders the S1/S2 border cleavage site inaccessible.

**Figure 5:**
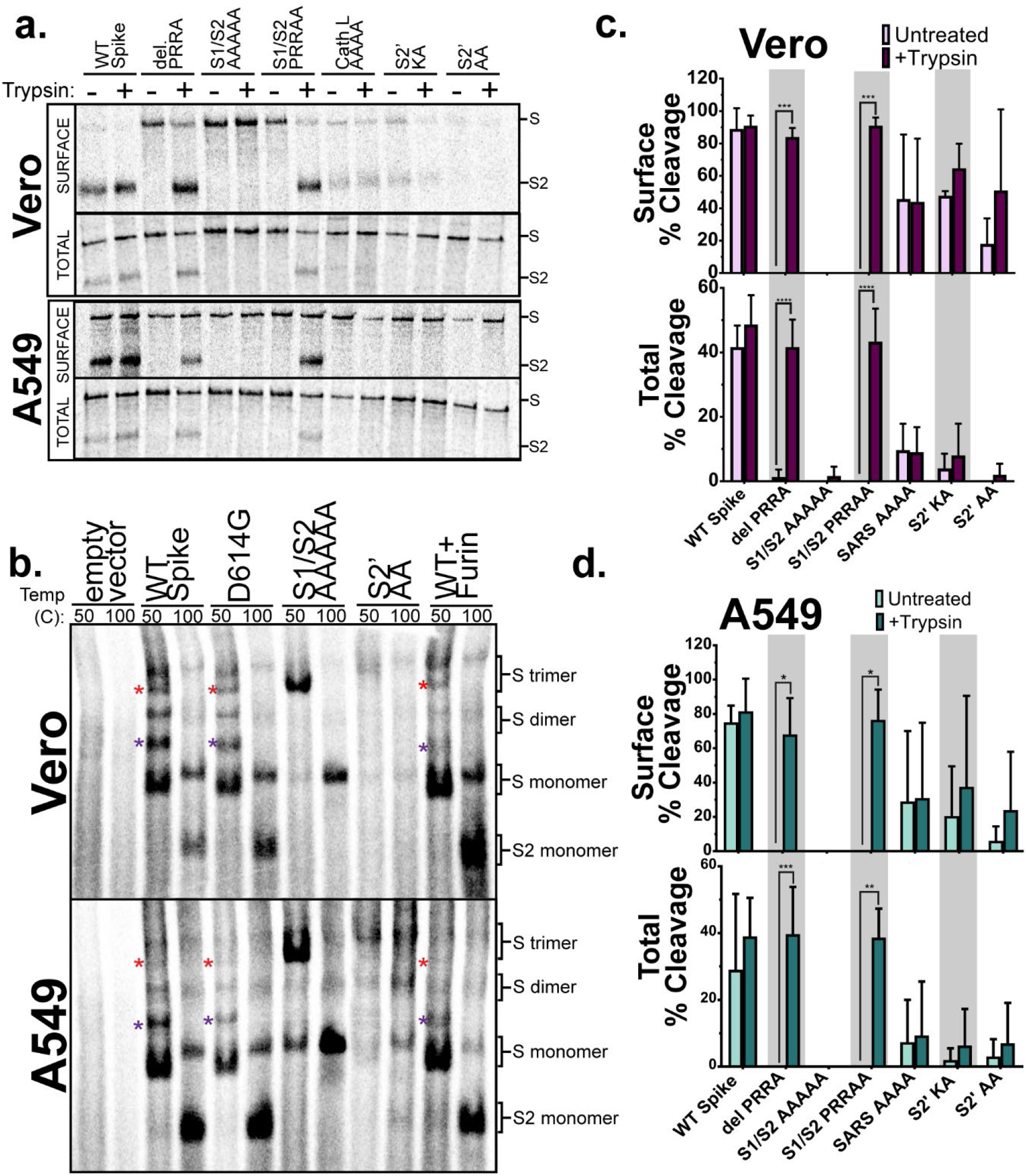
Mutations at downstream potential cleavage sites render the S1/S2 border cleavage site less accessible to proteases. a) Veros or A549s expressing wt S or S cleavage mutants were metabolically labeled for 6 hours. Surface proteins were biotinylated, and samples were either treated for 10 minutes with TPCK-Trypsin or left as untreated controls (as indicated). b) Veros or A549s expressing indicated proteins were metabolically labeled for 6 hours. Samples were treated at the indicated temperatures before separation on a nonreducing SDS-PAGE. Oligomers are labeled on the right based on size, and colored * represents potential intermediate species (n=3). Using band densitometry to quantify the bands in (a), percent cleavage was measured in (c) Vero and (d) A549 cells for both the surface (top graphs) and total (bottom graphs) populations (average ± SD for 3 independent experiments). Significance was determined by two-way ANOVA, *: p<0.05, **: p<0.01, ***: p<0.0005, ****: p<0.0001.

CoV S proteins associate as homo-trimers shortly after synthesis and remain in this trimeric form throughout the fusion cascade [12, 15]. To determine if proteolytic processing affects the stability of S trimer association, Veros or A549s transfected with wt S or mutants D614G, S1/S2 AAAAA, S2’AA, or wt S plus additional furin, were metabolically labeled. After lysis and immunoprecipitation, samples were then treated at 50°C or 100°C prior to separation on non-reducing SDS-PAGE. When wt S was incubated at 50°C prior to separation, species that correspond to a full-length S monomer, dimer, and trimer were observed (Fig. 5c). Interestingly, species that fall in between sizes corresponding to a monomer, dimer, and trimer (Fig. 5c, red and purple *) were also observed. These intermediate species may be the result of dimers or trimers made up of a mixture of full-length S protomers and cleaved S protomers. When wt S was incubated at 100°C prior to separation, bands corresponding only to full length S monomer, dimer, trimers, and cleaved S2 monomers were apparent. Similar results were also observed in D614G samples, suggesting that species containing cleaved protomers may be less stable. Consistent with this data, the S1/S2 AAAAA mutant, which cannot undergo cleavage at the S1/S2 border site, migrated primarily as a trimeric species after 50°C incubation, with little monomer or dimer observed. Additionally, when wt S was co-expressed with furin (shown to increase S cleavage in Fig. 1e and 1f), the predominant observed species was monomeric, after both 50°C and 100°C incubation. Overall, these results suggest that cleavage at the S1/S2 border alters the stability of S trimeric association.

### Furin or furin-like proteases in bat cells can cleave the S1/S2 border of SARS-CoV-2 Spike

*Rhinolophus affinis* horseshoe bats have been identified as the likely reservoir species for the novel SARS-CoV-2 [25]. To understand the proteolytic processing, expression, and stability of CoV-2 S in a cell line closely related to its reservoir host, we utilized *Pteropus alecto* fetus (pt. fetus) or lung (pt. lung) cells [64] that have a furin enzyme with ∼90% sequence homology to bats in the *Rhinolopus* family. Our previous studies on paramyxovirus virus fusion protein cleavage have shown that efficient furin and cathepsin cleavage occurs in these cells, although the furin cleavage occurs with delayed kinetics compared to Veros or A549s [65].

Surface biotinylation demonstrated that wt S and the del. PRRA mutant were readily expressed at the surface at similar levels in both cell lines, with cleavage at the S1/S2 border only observed for wt S and not for the del. PRRA mutant (Fig. 6a and 6b). Pulse-chase analysis showed that S expressed in both pt. lung and pt. fetus cells was cleaved at the S1/S2 border by one hour, with cleavage extent reaching approximately 40% at eight-hours, and 60% at 24 hours (Fig. 6c and 6d). Thus, furin or other proteases in *P*.*alecto* cells are able to process S, although this processing occurred more slowly than in other mammalian cell lines (compare to Fig. 1b). Interestingly, some cleavage was also observed in both pt. lung and pt. fetus cells for the del. PRRA mutation (Fig. 6c and 6d). Additionally, the wt S and del. PRRA mutant were slightly less stable in the *P. alecto* cells, demonstrating about 30-50% protein remaining at eight hours, and about 20% at 24 hours (Fig. 6e). In contrast, previously used mammalian cells lines showed 60-90% of wt S remained at eight hours, with 30-50% at 24 hours of chase (Fig. 1c).

**Figure 6:**
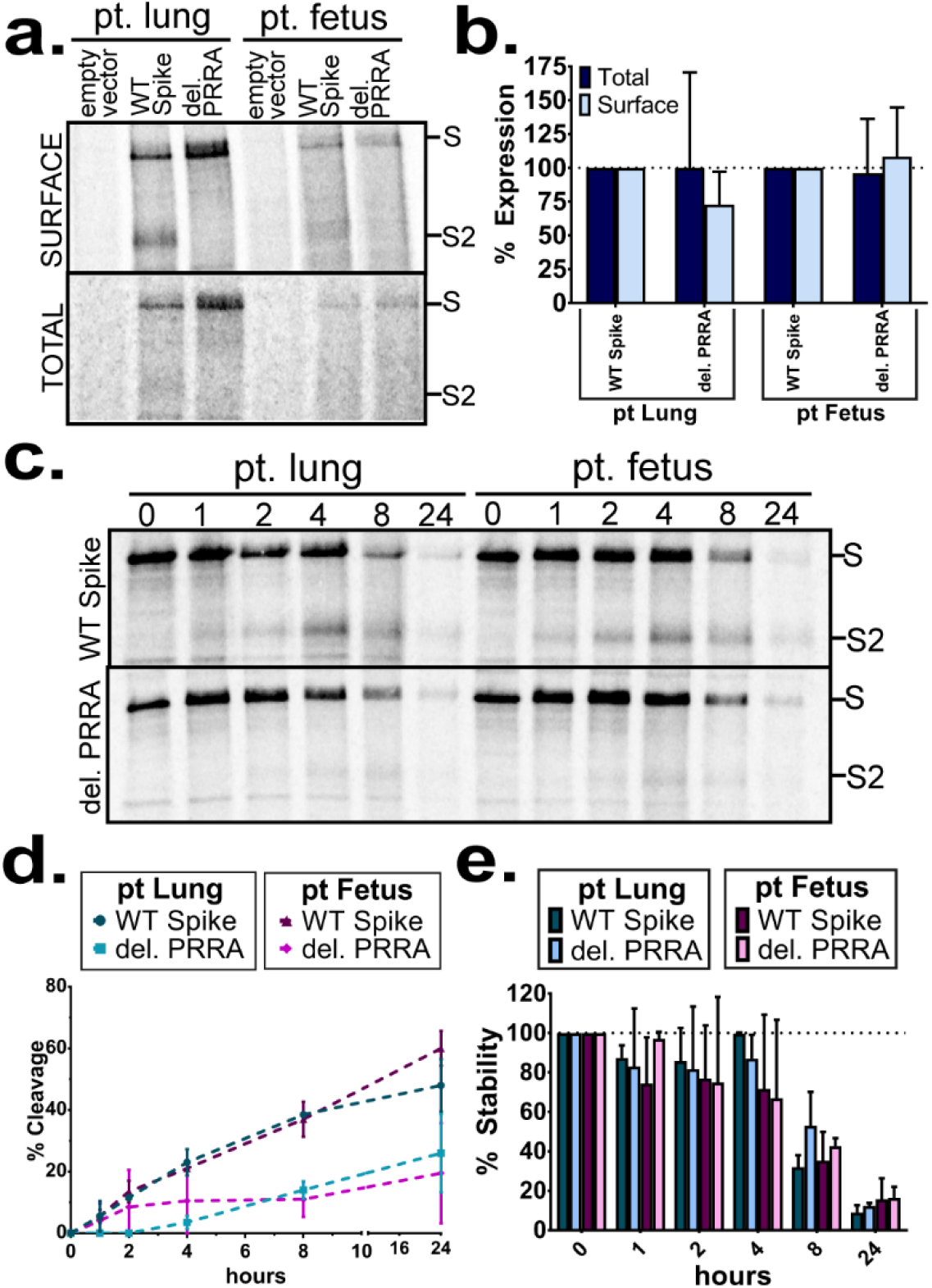
Furin or furin-like proteases in pteropus bat cells can cleave the S1/S2 border site of SARS-CoV-2 Spike. a) Surface biotinylation was performed on pteropus lung and pteropus fetus cells transfected with wt S or the del. PRRA mutant. b) Surface or total protein expression levels were quantified using band densitometry and normalized to wt S levels. c) pt. lung and pt. fetus cells were transfected with wt S or del. PRRA mutant, metabolically labeled for one hour, and chased for the times indicated. Again, using band densitometry to quantify bands results were expressed as (d) protein cleavage and (e) protein stability over time. (b,d,e average ± SD for 3 independent experiments)

## Discussion

In this study, we present a detailed characterization of the cleavage patterns, protein stability, and cell-cell fusion function of the SARS-CoV-2 S protein, as well as analysis of mutations within the S2 subunit that may affect these important protein properties. Consistent with recently published work [26, 35, 47, 50, 51, 66], our analysis confirms that S is readily cleaved at the S1/S2 border in a variety of mammalian cell lines. Additionally, we show for the first time, that cleavage occurs in a bat cell line similar to the SARS-CoV-2 reservoir species. While cleavage appears to be primarily carried about by the cellular protease furin, the sequence at this border does have the ability to be cleaved by other members of the pro-protein convertase family when furin is not present [47], and this likely accounts for the small amount of cleavage we observed in furin-negative LoVo cells.

Additionally, we carefully assessed the role different proteases play in cell-cell fusion, finding that furin increases cell-cell fusion when present in the same cell as S, and TMPRSS2 increases cell-cell fusion when present in a target cell, consistent with previous studies [37, 46]. Interestingly, when cell-cell fusion assays were performed using A549 cells as the effector cell (Fig. 2c), high background fusion levels were observed. This could be due to high endogenous levels of TMPRSS2 in this cell line compared to Veros, that were ultimately used for this experiment (Fig. S2c). High TMPRSS2 expression or exogenous treatment with trypsin has been shown to restore cell-cell fusion in low ACE2 receptor expression environments for SARS-CoV S [67, 68].

It is also worth noting that the presence of TMPRSS2 in the target (BSR/T7) cells also appears to process hACE2 (Fig. S2c, [40]). Therefore, we cannot exclude the possibility that the increase in fusion observed when TMPRSS2 is present in these cells is due to an effect on hACE2. In addition to the effect of proteases on cell-cell fusion, we also assessed the effect of Neuropilin-1, which has been suggested to be a co-receptor for SARS-CoV-2 viral entry and may be key for SARS-CoV-2 infiltration of the neuronal network [53-55]. We show that the presence of Neuropilin-1 with hACE2 in target cells does not impact S mediated cell-cell fusion (Fig. 2e). Additionally, co-expressing Neuropilin-1 with S in effector cells did not have an inhibitory effect on cell-cell fusion. While reports suggest Neuropilin-1 plays a role in viral entry of SARS-CoV-2, this indicates it does not play a significant role in S cell-cell fusion in our assay, although this was not investigated in neuronal cells.

The viral entry and cell-cell fusion pathways of SARS-CoV, MERS-CoV, and SARS-CoV-2 have several noteworthy commonalities, but do have marked differences. They all share the ability to facilitate entry through endosomal pathways, with S proteolytic activation mediated by endosomal/lysosomal proteases [10, 19, 35, 37-39, 69-72]. Additionally, they all can utilize cell surface (such as TMPRSS2) or extracellular proteases (trypsin) for S activation and subsequent viral entry [10, 37, 38, 47, 67, 72-78]. SARS-CoV-2 and MERS-CoV S differ from SARS-CoV S in that their S1/S2 border harbors a canonical furin cleavage motif [26, 27, 32], resulting in S pre-activation by furin during synthesis and cellular trafficking, prior to reaching the cell surface or being incorporated into viral particles [19, 35, 37, 39, 75]. This pre-activation by furin likely enhances the ability of SARS-CoV-2 and MERS-CoV S to participate in cell-cell mediated fusion without over-expression of cell surface or extracellular proteases [37, 46]. Addition of this cleavage sequence in SARS-CoV S allows SARS-S to facilitate cell-cell fusion without exogenous proteases [37, 79]. We show an increase in both syncytia formation and luciferase reporter gene assay fusion when cleavage at the S1/S2 border is enhanced by overexpression of furin (Fig. 2b and 2c), confirming that furin cleavage of SARS-CoV-2 S plays a critical role in cell-cell fusion. Interestingly, furin cleavage is not required for SARS-CoV-2 infection [10, 35, 37, 47], although removal of the site or inhibition of furin does appear to attenuate the virus [35, 39, 47] and reduce cellular tropism [46].

The presence of a furin consensus sequence is not only a marked difference between SARS-CoV and SARS-CoV-2, but it is also one of the differences between SARS-CoV-2 and a similar CoV circulating in a bat population [25]. Analysis of SARS-CoV-2 wt S in *P. alecto* cells demonstrates that this motif can be recognized and cleaved by furin in these cells (Fig. 6c and 6d), although the kinetics of this cleavage are noticeably slower than in other mammalian cell lines (compare to Fig. 1b). Previous work has shown that the fusion proteins of Hendra virus, processed by cathepsins, and parainfluenza virus 5, processed by furin, are also cleaved in *P. alecto* cells [65]. Pulse-chase analysis in this prior study demonstrated an increase in processing kinetics, although this kinetic difference can be accounted for by differences in protease expression levels between different bat cell lines (pt. kidney cells in [65], and pt. lung and pt. fetus cells in our work), suggesting there may be cellular differences in protein trafficking or furin activity. Intriguingly, a CoV-2 S mutant with a deletion of the inserted PRRA residues still demonstrated some cleavage in both utilized bat cell lines (Fig. 6c and 6d), while not showing any in Veros or A549s (Fig. 3c and 3d). Earlier work on MERS-CoV S showed that furin or other proprotein convertases in bat cells can process MERS S S1/S2 border without the presence of a canonical recognition motif [80]. Taken together, these results suggest that mutations in circulating bat CoVs that allow for human protease recognition at critical cleavage sites may be an important factor for zoonotic transmission of several CoVs.

Two other potential cleavage sites have been identified in work with other CoVs. The S2’ site is essential for both SARS and MERS infection [12, 32, 81-83] while a cath L activated site play a critical role for SARS-CoV S [13, 20, 84, 85]. Interestingly, mutations made at the S2’ site of SARS-CoV-2 S significantly reduce S1/S2 border cleavage, both in our study and others (Fig. 3b-d, [46, 86]), even though the sites are distal from each other. A similar reduction in cleavage is observed when the conserved cathepsin site is mutated (Fig. 3b-d). Our analysis of the published structures [3, 4, 87, 88] indicates that a full alanine mutation of this site may simply collapse the exposed S1/S2 loop. Our finding that exogenous trypsin treatment of cells expressing the S2’ or cathepsin site mutants does not restore cleavage at the S1/S2 border (Fig. 5a and 5b) suggests that these mutations result in proteins with altered furin loop structure [87], rendering it inaccessible. However, these mutants are still synthesized and trafficked to the surface despite not being cleaved (Fig. 3e-g), thus this change in conformation is unlikely to have drastically misfolded the protein. These results suggest that there may be a dynamic interaction between the S1/S2 border and S2’ cleavage sites in SARS-CoV-2 S needed to facilitate viral entry and cell-cell fusion. This dynamic control could also be regulated by S receptor binding exposing cryptic protease sites, although studies analyzing this in SARS and MERS S conflict on this topic [19, 70, 77, 89, 90].

We also assessed the effect on protein stability, cleavage, and cell-cell fusion function of a series of mutations in other regions of S. The D614G mutation emerged during 2020, and is now found in most circulating strains globally [48]. D614G has been shown to increase S incorporation into viral particles [91], increase receptor binding [92, 93], and reduce S1 subunit shedding and particle infectivity [94]. Importantly, the D614G mutant shifts S to favor a “heads up” conformation of the receptor binding domain [93, 95, 96]. In our study, the D614G mutation did not impact the cell-cell fusion function (Fig. 4f), expression, or stability of the protein (Fig. 4d/e, and Fig. S1), consistent with one previous study [86]. Our fusion results however conflict with two previous studies that demonstrated D614G increases cell-cell fusion, measured by cell depletion in flow cytometry [92], and syncytia formation in 293T and Hela cells stably expressing hACE2 [97]. These discrepancies may be due to differences in experimental conditions or cell types utilized. We are, however, the first to date to utilize a luciferase reporter gene assay to quantitate cell-cell fusion of a D614G S mutant. Using this assay, we also show that mutations found at two other residues (discovered in small, non-dominant population subsets [49]) alter the cell-cell fusion activity of S (Fig. 4f) without changing the overall protein expression or stability levels (Fig. 4d-e, Fig. S1d). Mutations at D839, a residue within the internal fusion peptide, to the polar amino acids, tyrosine or asparagine, significantly reduce fusion. Interestingly, a mutation at this residue that conserves the negative charge, D839E, has no effect on fusion activity. The negative charge at this residue may play a role in the regulation of S mediated fusion due to its location in the internal fusion peptide. Alternatively, this residue is in close proximity to C840, which may participate in a disulfide bond, so mutations at D839 may disrupt this disulfide bond, destabilizing the protein and changing fusion activity. Additionally, mutation of residue P1263 to a leucine significantly increases S mediated cell-cell fusion, suggesting that residues in the cytoplasmic tail may play a role in the S-promoted cell-cell fusion process. Notably, a study that removed the entire S cytoplasmic tail still observed syncytia formation at levels similar to wt S [86], indicating that regulation by the cytoplasmic tail may be complex or that the role of the cytoplasmic tail in fusion is not regulation, but interaction with cellular host factors [98].

In this work, we also provide critical insight into the kinetics of protein cleavage and overall stability of CoV-2 S. S protein processing at the S1/S2 border occurs within two hours of synthesis (Fig. 1a and 1b; one hour of label, one hour of chase) in several mammalian cell lines (Vero, MEF, A549), and continues to increase over time, reaching 60-80% protein cleavage by eight hours of chase time, depending on the cell type. Overexpression of furin increased the efficiency of S1/S2 border cleavage (Fig. 1d-f), and this increase in cleavage may account for the increase in cell-cell fusion observed when furin is co-expressed with S (Fig. 2a-c, [37, 46]). Additionally, we show that transiently transfected S is stable in several mammalian cells for 4-5 hours post-protein synthesis with demonstrable turnover after this point, (Fig. 1c, Fig. S1). This protein turnover is similar to turnover rates seen in PIV5 fusion protein, also activated by cellular furin [99], and slightly slower turnover than Hendra fusion protein, activated by cellular cathepsins [100, 101]. Over-expression of cellular proteases that may process S did not affect these protein turnover rates. Interestingly, analysis of S in non-reducing conditions found that cleavage of the S1/S2 border appears to destabilize trimeric interactions (Fig. 5b). In these non-reducing conditions, no differences were observed in oligomeric stability between wt S and the D614G S mutations, despite the D614G favoring a ‘heads up’ conformation [93, 95, 96] and Vero cells having sufficient levels of endogenous ACE2 to facilitate syncytia formation (Fig. S2c), suggesting that changes in receptor binding do not alter overall protein trimeric association. Notably, in these non-reducing conditions after a 50°C treatment for wt S, the D614G mutant, and wt S+furin, bands between monomer, dimer, and trimer species are observed (Fig. 5b, indicated with *). These intermediate species are not observed after treatment at 100°C. These may represent protein oligomers that are not identically cleaved and are therefore partially destabilized, a phenomenon proposed for MERS-CoV S [32], and murine hepatitis virus CoV S, [102]. Protein oligomers with differential proteolytic processing may also account for the small population of un-cleaved protein we observed at the cell surface in our experiments (Fig. 3e, Fig. 4d, Fig. 5a, and Fig. 6a).

Through biochemical and cell biological analysis of the SARS-CoV-2 S protein, we have provided important observations about the stability, proteolytic processing, and requirements for cell-cell fusion of this highly sought-after therapeutic target. This information may be helpful in directing treatments that inhibit S protein fusion, or for discerning methods to stabilize CoV-2 S in therapeutic development. Additional studies are needed to understand the potential interplay between S cleavage sites and how that may contribute to S protein function, as well as to further investigate spike S2 subunit regions that are critical for protein function.

## Experimental procedures

### Cell lines and culture

Vero (ATCC), BSR T7/5 cells (provided by Karl-Klaus Conzelmann, Pettenkofer Institut), mouse embryonic fibroblasts (MEFs) from cathepsin L knockout mice (Cath L-MEFs) (a gift from Terence Dermody, University of Pittsburgh), and *P. alecto* bat cells harvested from fetus (pt. fetus) and lung (pt. lung) (a gift from Linfa Wang, Duke-NUS) [64] were all maintained in Dulbecco’s modified Eagle’s medium (DMEM, GE Healthcare), with 10% fetal bovine serum (FBS) and 1% penicillin/streptomycin. Every third passage, 0.5mg/ml of G-418 (Invitrogen) was added to the culture media of BSR T7/5 cells to select for the expression of the T7 polymerase. A549 and human colon carcinoma LoVo cells (both purchased from ATCC) were cultured in F12 Kaighns Modification media (GE Healthcare) with 10% FBS and 1% penicillin/streptomycin.

### Plasmids, Antibodies, and Mutagenesis

pCAGGS-SARS-CoV-2 spike was obtained from BEI Resources. pcDNA3.1(+)-hACE2 and pcDNA3.1(+)-TMPRSS2 were provided by Gaya Amarasinghe (Washington University). Human Neuropilin-1 was expressed with an exogenous PTPα signal sequence from the pLEXm vector (from Craig Vander Kooi, University of Kentucky). SARS-CoV-2 S was subcloned into pUC57 and all S mutants were created in pUC57 using the QuikChange site-directed mutagenesis kit (Strategene) with primers purchased from Eurofins. Constructs were then subcloned back into the pCAGGS expression vector. Other plasmids utilized include pSG5-Cathepsin L (from Terence Dermody, University of Pittsburgh), pCAGGS-furin (Promega), and T7 promoted-luciferase (Promega). Antibodies anti-SARS spike glycoprotein (ab252690) and anti-hACE2 (ab15348) were purchased from Abcam, and anti-TMPRSS2 (H-4) was purchased from Santa Cruz Biotechnology, Inc.

### Gel electrophoresis and western blotting

Proteins were separated on a 10% sodium dodecyl sulphate-polyacrylamide gel electrophoresis (SDS-PAGE). For western blot analysis, proteins were transferred to a polyvinylidene difluoride (PVDF) membrane (Fisher Scientific) at 60V for 100 minutes. After blocking with 5% milk in tris-buffered saline + Tween-20 (tTBS) for 1 hour, membranes were incubated with respective antibodies (anti-SARS S 1:5000 dilution, anti-TMPRSS2 1:1000 dilution, anti-hACE2 1:1000 dilution) at 4°C overnight. Membranes were then washed with tTBS and incubated with (Li-Cor) secondary antibodies at 1:10000 dilution in 5% milk solution for 1 hour. Membranes were washed again with tTBS and diH_2_O, before being imaged on the Odyssey Image Analyzer (Li-Cor).

### Syncytia Assay

Cells (Vero or A549s) in 6 well plates were transiently transfected with 2μg of either wild-type or mutant SARS-CoV-2 S protein plasmid with Lipofectamine 3000 (Invitrogen) at a ratio of 1:2:2 DNA: P3000: Lipofectamine 3000. For experiments with the addition of proteases, the total DNA transfected was kept constant at 2μg, in those cases we used 1μg of S and 1μg of the indicated protease. Syncytia formation was imaged at 24 and 48 hours post transfection on a Nikon Ti2 at 20X magnification.

### Luciferase Reporter Gene Assay

Effector cells (Vero or A549s) were plated in 12-well plates at 70-90% confluency and transfected with 1μg of total DNA (0.4μg of a T7 promoted luciferase plasmid, 0.6μg of wild-type (wt) or mutant S protein or S protein with additional proteases). At the same time BSR cells (constitutively expressing a T7 promoter) seeded in 6-well plates were transfected with 2μg either empty pCAGGS or pcDNA3.1(+)-hACE2. Eighteen to twenty-four hours post transfection BSR cells were lifted using trypsin, centrifuged for five minutes at 1500 rpm, resuspended in normal DMEM+10% FBS, and overlaid onto the S expressing cells at a ratio of 1:1. Overlaid samples were then incubated at 37°C for 9 hours (or as described in the text). Samples were lysed in 100μL of Reporter Gene Lysis buffer (Promega) and frozen overnight. Plates were then scraped on ice, lysates were vortexed for 10 seconds, centrifuged at 13,000 rpm for 1 minute at 4°C, and 20μL of the supernatant was added to an opaque 96 well plate. Luciferase activity was measured on a SpectraMax iD3 (Molecular Devices) using a Luciferase Assay System (Promega). Background values were subtracted (empty pCAGGS in BSRs and effector cells) and luciferase activity was expressed as a percentage of wt S (effector cells) and hACE2 (BSR cells).

### Surface Biotinylation

Two μg of wt or mutant S protein was transfected into Vero or A549 cells using the Lipofectamine 3000 system (Invitrogen; ratios described above). Eighteen to twenty-four hours post transfection, cells were starved in Cys^-^/Met^-^media (Gibco) for 45 minutes, and metabolically labelled for six hours using 50μCi of S^35^ (PerkinElmer) incorporated into Cys and Met (S^35^ Cys/Met). After the label, cells were washed once with PBS (pH 8) and incubated with 1 mg/ml of EZ-link Sulfo-NHS-biotin (Thermo Fisher) in PBS (pH 8) at 4°C for 35 minutes, and then at room temperature for 15 minutes. Next the cells were lysed in 500μl of RIPA buffer (100 mM Tris-HCl [pH 7.4], 0.1% SDS, 1% Triton X-100, 1% deoxycholic acid) containing 150 mM NaCl, protease inhibitors (1 U aprotinin, 1mM PMSF, [both from Sigma-Aldrich]), 5 mM iodoacetamide, and cOmplete EDTA-free Protease Inhibitor Cocktail Tablets (all from Sigma-Aldrich). Cell lysates were centrifuged at 55,000 rpm for 10 minutes, and the supernatant was incubated with anti-SARS S polyclonal antibody at 4°C for three hours. Following incubation, Protein A conjugated to Sepharose beads (Cytiva) were added to the samples, and incubated at 4°C for an additional 30 minutes. Post-incubation samples were washed two times with each RIPA Buffer+0.3M NaCl, RIPA Buffer+0.15M NaCl, and SDS-Wash II buffer (50mM Tris-HCl [pH 7.4], 150mM NaCl, and 2.5mM EDTA). After buffer aspiration and addition of 10% SDS, samples were boiled for 10 minutes. The supernatant was removed to a separate tube. 15μl of supernatant was removed and added to an equal portion of 2X SDS loading buffer and labeled “TOTAL”. Biotinylation buffer (20 mM Tris [pH 8], 150mM NaCl, 5mM EDTA, 1% Triton X-100, and 0.2% BSA) and Streptavidin conjugated beads were added to the remaining supernatant, and this was incubated at 4°C for one hour. Samples were again washed as described above and 2X SDS loading buffer was added following the washes. Samples were boiled for 15 minutes and run on a 10% SDS-PAGE gel. Gels were dried and exposed on a phosphoscreen for two to four days, then visualized using a Typhoon Imaging System (GE Healthcare). Bands were quantified using band densitometry using the ImageQuant software (GE Healthcare).

### Time Course Immunoprecipitation

2μg of wt or mutant S was transfected into Vero or A549 cells using the Lipofectamine 3000 system (Invitrogen; ratios described above). Eighteen to twenty-four hours post transfection, cells were starved in Cys^-^/Met^-^media (Gibco) for 45 minutes, and metabolically label for one hour using 50μCi of S^35^ Cys/Met. After the one-hour label, cells were washed once with PBS and normal DMEM + 10% FBS was added for indicated times. Cells were then lysed in 500μl of RIPA lysis buffer. Anti-SARS S polyclonal antibodies were used to immunoprecipitate the CoV-2 S protein as previously described and the protein was analyzed on a 10% SDS-PAGE gel. Gels were dried and exposed on a phosphoscreen for 2-4 days and visualized using a Typhoon Imaging System (GE Healthcare). Bands were quantified using band densitometry using the ImageQuant software (GE Healthcare).

### Non-reducing Gel Electrophoresis

Two μg of wild-type or mutant S was transfected into Vero or A549 cells using the Lipofectamine 3000 system (Invitrogen; ratios described above). Eighteen to twenty-four hours post transfection, cells were starved in Cys^-^/Met^-^media (Gibco) for 45 minutes, and metabolically labeled for six hours using 50μCi of S^35^ Cys/Met. Lysed cells were immunoprecipitated as described above, however after the washing steps, 30μl of 2X SDS loading buffer without dithiothreitol (DTT) was added to each sample. Samples were treated at 50°C or 100°C, as indicated, for 20 minutes and analyzed on a 3.5% acrylamide gel under non-reducing conditions. The gel was dried, exposed, and imaged as described for surface biotinylation.

### Immunofluorescence experiments

Sub-confluent cells on coverslips in 6 well plates were transfected with 2μg of DNA using the Lipofectamine 3000 transfection system (Invitrogen). Eighteen to twenty-four hours post transfection cells were fixed with 4% PFA for 15 minutes at room temperature. Cells were permeabilized in a solution of 1% Triton X-100 in PBS+0.02% Sodium Azide (PBSN) for 15 minutes at 4°C. After permeabilization, coverslips were moved to a humidity chamber and blocked with 1% normal goat serum (NGS) in PBSN for 1 hour at 4°C. Cells were labeled with anti-SARS S antibody (1:2000 dilution) in blocking buffer overnight at 4°C or for three to five hours at room temperature. Samples were washed with PBSN+0.01% Tween-20 seven times and incubated for 1 hour at 4°C with goat anti-rabbit FITC (1:2000 dilution). Samples were again washed with PBSN+0.01% Tween seven times and mounted onto slides using Vectashield mounting media (Vector Laboratories). Slides were imaged on an Axiovert 200M (Zeiss) at 63x magnification using Metamorph to collect Z-stacks and processed using Nikon NIS Elements.

### Statistical analysis

Statistical analysis was performed using Prism 7 for Windows (GraphPad). A *p* value of <0.05 was considered statistically significant. Multiple comparison tests were generated using one-way or two-way analysis of variance (ANOVA) with Dunnett’s multiple comparison test. *: p<0.05, **: p<0.01, ***: p<0.0005, ****: p<0.0001

### Data availability

The datasets generated during and/or analyzed during the current study are available upon request from the corresponding author, Rebecca Dutch, on reasonable request.

## Acknowledgments

We would like to thank Craig Vander Kooi (University of Kentucky) for providing the Neuropilin-1 expression plasmid and for providing structural insight. We would also like to thank Gaya K. Amarasinghe (Washington University School of Medicine) for providing the TMPRSS2 and hACE2 expression plasmids, and providing feedback regarding experimental design.

## Funding and additional information

Financial support was provided by the CCTS CURE Alliance pilot award from the University of Kentucky, NIAID grant R01AI051517 to R.E.D. and NIAID grant R01AI140758 to R.E.D and D.W. L.

## Conflict of interest

The authors declare that they have no conflicts of interest with the contents of this article.

